# Linear gain effect of theta-burst and 15 Hz rTMS on intra-cortically measured neuronal responses to oriented grating visual stimuli

**DOI:** 10.1101/2020.03.09.983932

**Authors:** Ajay Venkateswaran, Ze-Shan Yao, Martin Y. Villeneuve, Amir Shmuel

## Abstract

Transcranial magnetic stimulation (TMS) involves the application of time-pulsed magnetic fields to cortical tissue through a coil positioned near the head. TMS is widely used for studying the mechanisms underlying perception and behaviour and is considered a potential therapeutic technique for various conditions. However, the application of TMS has been hindered by the lack of understanding of its mechanism of action. Here we studied the effects of three repetitive TMS (rTMS) paradigms on intra-cortical neuronal responses to oriented gratings in cat area 18. Each of stimulation protocols, including continuous theta-burst, intermittent theta-burst, and 15 Hz rTMS, consisted of 600 pulses. Application of continuous theta-burst and 15 Hz rTMS suppressed the action potential response to oriented grating stimuli for 1.5-6 minutes. In contrast, application of intermittent theta-burst stimulation was associated with enhancement of the action potential response ∼15 minutes following TMS. Neuronal assemblies that were more responsive before the application of rTMS, were affected more than neurons that were less responsive before rTMS. In spite of these changes, the preferred orientation, the orientation tuning of the multi-unit activity and the spatial pattern of the responses recorded from assemblies neurons remained unaffected. Our findings demonstrate that rTMS does not modify the functional selectivities of ensembles of neurons; rather, it has a linear gain effect on their responses.

## Introduction

Transcranial magnetic stimulation (TMS) involves the application of time-pulsed magnetic fields to cortical tissue through a coil positioned near the head. Since its inception (Barker et al., 1985a), TMS has been considered as a potential therapeutic technique for conditions of recovery from stroke (Ernst, 1990; Khedr et al., 2009; Leong, 2009; Takeuchi et al., 2005), psychiatric disorders such as depression (Berlim et al., 2012; Berlim et al., 2013), schizophrenia (Prikryl and Kucerova, 2013; Rajji et al., 2013) and others. It has been used in pain research and in studies involving conduction pathways of the central and the peripheral nervous systems (Awad et al., 2013; Massé-Alarie et al., 2013; Schabrun et al., 2013). TMS has also been commonly used to investigate the relationship between brain and behavior (Rossini et al., 2010). However, the application of TMS has been hindered by the lack of understanding of its mechanism of action, and therewith, the lack of tools for predicting its effect on cortical processing (Wasserman and Zimmerman, 2012). Therefore, most of the concepts on the rTMS’ mechanisms of action have been speculative.

Various TMS delivery paradigms cause different effects that last for different durations. The effect of single-pulse TMS lasts, at most, a few tens of milliseconds (Rossi et al., 2009; Pascual-Leone, 2002). Therefore, it is generally used as an ‘instantaneous’ paradigm in which TMS is delivered during the performance of a task. Repetitive TMS (rTMS), on the other hand, is used both as an ‘instantaneous’ and ‘prolonged’ paradigm, since the duration of its effect is on the order of seconds or minutes, outlasting the application of the pulses (Rossi et al., 2009; Pascual-Leone, 2002). Based upon the frequency of repetition of pulses, rTMS can be further classified into low frequency (<= 1Hz) and high frequency (>1 Hz) rTMS. Low and high-frequency rTMS paradigms are known to induce significant effects on cortical excitability and plasticity (Valero-Cabre et al., 2007; Gersner et al., 2011; Siebner, 2010; Vlachos et al., 2012). Low frequency rTMS has been shown to decrease cortical excitability and hence increase seizure threshold (Tergau et al., 1999; Joo et al., 2007; Prikryl and Kucerova, 2005). The converse may be true for high-frequency rTMS, which increases cortical excitability (Berardelli et al., 1998), and therefore may be prone to induce seizures.

Several high-frequency TMS stimulation protocols have been used. Our interest lies in a particular rTMS protocol developed by (Huang et al., 2005). This technique conditions the brain with bursts of high-frequency rTMS. Each burst consists of 3 pulses at 50 Hz. These bursts can then further be repeated at a frequency of 5Hz (theta), 10 Hz (alpha) or 20 Hz (beta) to provide an rTMS paradigm which can be referred to as theta-, alpha-, or beta-burst stimulation, respectively (De Ridder et al., 2007).

Previous studies that combined continuous or burst rTMS paradigms with invasive electrophysiological recordings have been pursued in rodent (Benali et al., 2011; Levkovitz et al., 1999), feline (de Labra et al., 2007; Espinosa et al., 2007; Espinosa et al., 2011; Moliadze et al., 2005; Moliadze et al., 2003; Pasley et al., 2009), non-human primate (NHP) models (Baker et al., 1995; Dancause et al., 2006), and one human patient (Wagner et al., 2004a). The reported effects of TMS paradigms on cortical excitability and function are highly variable. Continuous theta burst (cTBS) applied to the motor cortex suppressed motor evoked potentials (MEPs) amplitudes – a peripheral measure of cortical excitability (Huang et al., 2005; Ishikawa et al., 2007). In contrast, Benali et al. (2011) reported an increase in the fundamental (in a train of three) somatosensory evoked potentials (SEPs) P_1_N_1_ amplitudes following cTBS conditioning. The opposite result was reported bt Zapallow et al. (2012). In their hands, the P_1_N_1_ amplitude is significantly suppressed for 5 to 9 minutes following cTBS. In addition, Ishikawa et al. (2007) and Zapallow et al. (2012) reported that cTBS suppressed a different SEPs component – P_25_N_33_ or effectively P_2_N_2_. Possibly explaining part of the variability, Noh et al. (2012) and (Vernet et al., 2013) showed that the effect of cTBS may depend on the site in which activity is measured by demonstrating that cTBS suppressed MEP amplitudes but had a longer lasting increase in EEG power amplitudes. In contrast to the reported effects of cTBS, intermittent theta burst (iTBS) shows consistent facilitatory effect on both MEPs and SEPs (Benali et al., 2011; Di Lazzaro et al., 2008; Huang et al., 2005). However, 15 Hz rTMS – a paradigm that delivers single pulses at the average frequency of the bursts used for cTBS – seems to cause variable effects. It has shown facilitatory effects on MEPs (Wu et al., 2000) and no effect on MEPs (Huang et al., 2005; Maeda et al., 2000a). Thus, previous reports on the effect of 15 Hz and cTBS on neurophysiological activity are inconsistent.

Not only the sign – suppression or facilitation – of the effect of rTMS on neurophysiological activity is not yet determined, their amplitudes too. Yet another important feature of the effect of rTMS paradigms is their time course. What’s the duration of the effect? How long after the application of rTMS do responses go back to their normal amplitudes?

In order to reconcile these inconsistent reports, we applied intracortical recordings of two more local measures of excitability, namely action potentials and gamma-band responses. To this end, we recorded neurophysiological responses to oriented grating stimuli from an array of electrodes implanted in cat area 18. We aimed to determine the effects of cTBS, iTBS, and 15 Hz rTMS paradigms on neuronal responses, including the sign – suppression or facilitation, amplitude of the response relative to the amplitude prior to rTMS, time-course and the spatial pattern of the responses.

## Materials and Methods

### Animals and anesthesia

All procedures were approved by the animal care committees of the Montreal Neurological Institute and McGill University and were carried out with great care according to the guidelines of the Canadian Council on Animal Care. Data were obtained from one male and six female cats, weighing 3.7 to 4.7 Kg. Twelve hemispheres (2 hemispheres from 5 cats and one hemisphere each from 2 cats) have been used to obtain 12 datasets for each TMS paradigm. Of the 12 datasets, 4 datasets were rejected: three due to unstable baseline activity (which makes it difficult to quantify the effect of TMS) and one due to seizure induction.

We performed acute experiments. Glycopyrrolate (0.01-0.05 mg/Kg) was administered intramuscularly (IM). Fifteen minutes later, we administered a cocktail of acepromazine (0.11 mg/Kg SC) and ketamine hydrochloride (20 mg/Kg, IM). The animals were then placed on a heating pad, and their body temperature was maintained within the range of 38.0°C-38.5°C). The cats were intubated and ventilated using a respiratory pump, with a mixture of 80% Medical Air, 20% Oxygen and isoflurane. Venous cut-downs were performed to insert cephalic vein catheters into both forelimbs. These two lines were subsequently used for all intravenous (IV) administrations.

The animals were carefully placed on a stereotaxic frame with their heads supported by ear bars coated with lidocaine hydrochloride gel. To prevent the dryness of the cornea, we applied contact lenses. Throughout the experiment, the animals’ heart rate (HR; measured with ECG), pulse oximetry (SPO_2_), end-tidal carbon dioxide (Et-CO_2_) and body temperature (T) were monitored and stabilized. Once all physiological parameters were stable, Gallamine Triethiodide (5 ml) was injected through one of the IV lines to induce paralysis, in order to prevent eye movements. All surgical wounds were infused with 2% lidocaine hydrochloride. Glycopyrrolate (IM every 7 hours at 0.01 mg/kg), Buprenorphine (IV every 12 hours at 8.5 µg/kg) and Dexamethasone (IV every 24 hours at 0.3 mg/kg) injections were administered to prevent tracheobronchial secretions, post-operative pain, and inflammation, respectively.

The cranium was exposed, and two holes were drilled for the insertion of small metal screws for recording the EEG. For optical imaging and for the subsequent implantation of an electrode array, a larger circular opening (approximately 18 mm in diameter) was made, centered on the midline at Horsley-Clark coordinate A4. A chamber was attached to the skull. In order to prevent edema, mannitol (250 mg/kg) was injected (IV) 30 minutes before performing duratomy. The dura mater above area 18 was resected. Artificial cerebrospinal fluid (aCSF) or agarose were used to protect the exposed cortex. At the end of the surgical procedures, gas anesthesia was switched from isoflurane to halothane (Sigma-Aldrich). An infusion of 6ml/kg/hr of 5% Dextrose in Lactated Ringer’s solution and 2% Gallamine Triethiodide (2 mg/Kg/hr) was administered throughout the duration of the experiment. During the imaging and recording sessions, the halothane level was kept at 0.7-1.0% and the heart rate was maintained at 160-180 BPM. The animal was mechanically ventilated at a rate of 25-40 strokes per minute and a volume of 10-15 ml/kg, maintaining the end-tidal CO2 within 32-38 mm Hg. The gas mixture was adjusted within the range of 100 % medical air to 80% medical air / 20% O2, to keep the oxygen saturation level at ≥ 94%.

To dilate the pupils, we administered 2.5% phenylephrine hydrochloride (Mydfrin®, Aventix Animal Health). The eyes were protected using contact lenses with zero power; they were focused on a tangent screen at a distance of 30 cm using external lenses with power determined by retinoscopy. The retinal vessels, blind-spot, and area-centralis of each eye were back-projected onto the screen.

### Visual Stimuli

To generate the visual stimuli, we used the Psychophysics toolbox (Brainard, 1997). The visual stimuli were displayed on a 21” LCD screen with a refresh rate of 60 Hz. The screen was positioned at a distance of 30 cm from the animal, subtending an angle of 30–55° in the visual field, contralateral to the hemisphere investigated. The animals were stimulated binocularly, using 50% contrast sinusoidal-wave gratings with a spatial frequency of 0.15 cycles per degree and a temporal frequency of 4 Hz.

To optimize the position of the electrode array, we used optical imaging to acquire hemodynamic responses to grating stimuli. Each presentation of a visual stimulus was associated with 12s-long data-acquisition, starting with 2s of presenting static gratings followed by 5s of moving grating and 5s of static gratings. To allow for the relaxation of activity-dependent vascular changes, each 12-second period of data-acquisition was followed by 8-second inter-stimulus interval (ISI). The stimulus was switched to the static grating of the next condition at the beginning of the ISI.

Each of 30 trials consisted of acquiring responses to 16 grating stimuli moving in 16 directions, spanning the orientation and direction spaces at a resolution of 22.5°. In addition, we acquired baseline optical images during a presentation of a blank image – an iso-luminant gray screen – of luminance equal to the average luminance of the gratings. All 17 conditions were presented once in each trial with a randomized order of presentation, averaging out any systematic effects of stimulus presentation order.

### Optical imaging and implantation of a multi-electrode array

Responses of neuronal assemblies were imaged optically and subsequently recorded with microelectrode arrays (MEAs). The surface of the cortex was imaged using a differential data acquisition system (VDAQ Imager 3001, Optical Imaging Ltd., Rehovot, Israel) equipped with a 12-bit bit-depth Pantera 1M60 camera (Teledyne Dalsa, Waterloo, Ontario, Canada) and a macro lens (Nikon, AF Micro Nikkor, 60 mm, 1:2.8 D). The camera was focused on the surface of the cortex, ensuring that the cortical blood vessel architecture was captured. Preferred orientation was mapped parallel to the surface of cortex (Grinvald et al., 1986; Shmuel and Grinvald, 2000). After obtaining the orientation map, an optimally responding region was selected for implanting a 10×10 electrode array (3.6 mm × 3.6 mm area, 400 μm separation between adjacent electrodes, and 1 mm electrode length; Blackrock micro-systems, Salt-Lake Cirt, UT) for electrophysiology recordings. The array was then inserted using a pneumatic inserter. Figure 1 shows an orientation map obtained during one of the experiments followed by the placement of the electrode array. After the insertion of the array, a laser beam was used to register the location of the array (Figure 1c). The array was then covered by a thin layer of Surgifoam soaked in artificial CSF (approximately 1 mm thickness in the soaked state) in order to replicate the CSF layer and keep the brain wet. The drilled out piece of bone was then placed above the array, now cushioned by a layer of wet Surgifoam.

**Figure 1.**
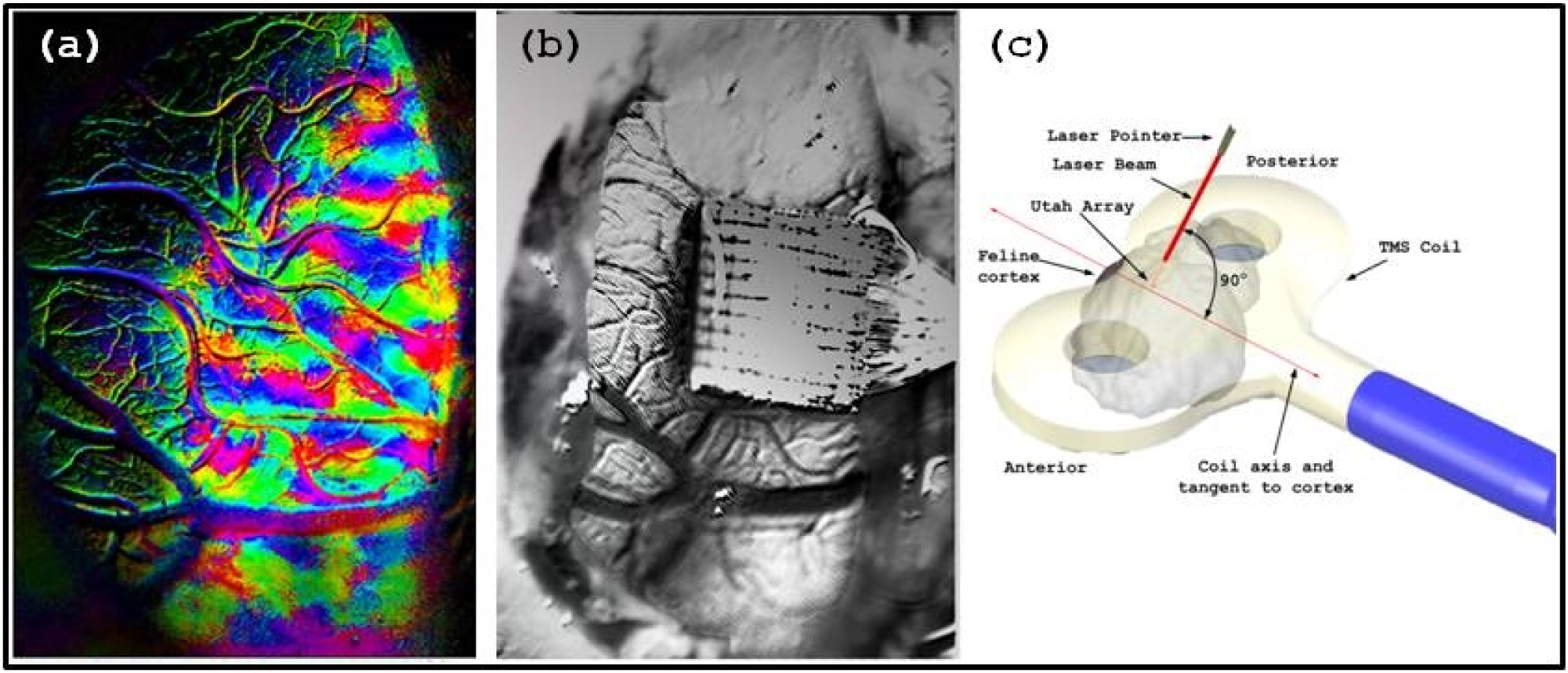
Mapping the area intended for array implantation followed by coil positioning. (a) Preferred orientation map obtained using optical imaging. (b) The position of the implanted array, determined based on the orientation map. (c) Localization of the TMS coil above the array, using a laser-guided pointer held perpendicularly above the array.

A 70 mm butterfly TMS coil was placed above the array approximately 10 mm from the cortical surface. Since the gyrus on which the array was inserted was directed in the anterior-posterior direction, the TMS coil was oriented tangentially to the cortex amd perpendicularly to the anterior-posterior axis. This caused the induced current profile to approximately align with the mediolateral axis (perpendicular to the direction of the gyrus). This orientation is important for inducing the maximum electric field that can be induced in a gyrus for a particular stimulation intensity (Thielscher et al., 2011).

### rTMS protocols and experimental paradigm

Trains of rTMS pulses of biphasic waveforms were generated by a Magstim Rapid TMS stimulator (Magstim, Whitland Dyfed, UK) and administered to cortical area 18 through the coil.

Three different rTMS paradigms were administered: a 15 Hz rTMS paradigm (a train of 15 Hz pulses for 40s amounting to a total of 600 pulses) and two theta-burst paradigms (three pulses of TMS at 50 Hz repeated at 200 ms intervals; Huang et al., 2005). We applied cTBS (a single train of TBS for 40 s amounting to 600 pulses) and iTBS (a 2s train of TBS repeated every 10 s for 190s amounting to 600 pulses). All TMS paradigms were delivered at 55% MSO, which equivalent to 100% RMT.

Figure 2 depicts our experimental paradigm associated with each of the 3 rTMS protocols. Each presentation of a visual stimulus consisted of 1s of a blank iso-luminant gray screen, followed by 1s of moving grating. Each 34s trial consisted of presenting 16 gratings stimuli spanning the orientation and direction space at a resolution of 22.5°, and 1 blank condition. The order of the directions in each trial was randomized. Seventy-five such trials were run for each TMS protocol. TMS was administered following the completion of 15 baseline trials, before the 16th trial started. Thus, the duration of one run was 8.5 minutes of pre-TMS and 34 minutes of post-TMS.

**Figure 2.**
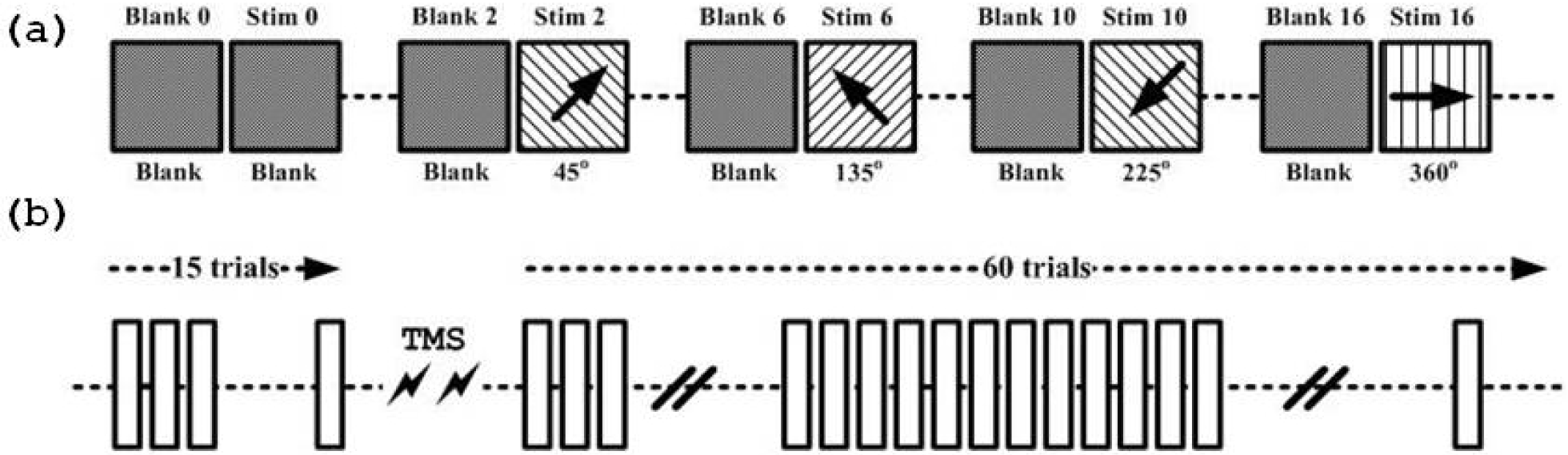
Experimental Paradigm. (a) Each trial consists of 16 conditions (8 orientations and 16 directions of motion orthogonal to orientation, spanning 360° in steps of 22.5°) and a blank condition. Each stimulus was presented for 1 s, with 1 s inter-stimulus interval during which a blank gray image was presented. (b) Each square block corresponds to one trial. We obtained data from fifteen trials before TMS, followed by the delivery of TMS and subsequently 60 trials following the application of TMS.

Throughout each run, extra-cellular neurophysiological signals were recorded continuously at a sampling rate of 24,414 Hz using a Tucker Davis Technology (Alachua, FL) data acquisition system. We recorded local field potentials and spiking activity.

### Data analysis

The results from each run consisted of 75 vectors ordered according to time of the acquisition, each of which containing the responses to 17 conditions during one trial. For analysis, we re-sampled the 75 vectors to 25 vectors by averaging 3 consecutive trials. Responses were averaged over the same orientation, maintaining the vector structure.

Statistical significance of the effect of TMS was tested using a t-test with p<0.05. Post-hoc false discovery rate (FDR) correction based on Bejamini and Yekutieli’s procedure was performed on all statistical tests. Statistically significant decreases and increases are represented by blue and red (*) markers, near the lower and upper y-axis extremities, respectively. Statistical significance of the correlation plots was obtained by transforming correlation coefficients to Fisher coefficients and performing a t-test on the Fischer coefficients. However, for presentation purposes, actual correlation coefficient values are presented. To evaluate the similarity between two distributions we applied a paired Kolmogorov-Smirnov test followed by a Chi-squared goodness-of-fit test.

### Sifting orientation-selective channels from other channels

Each evoked response to a visual stimulus was split into three segments: spontaneous segment (800 milliseconds baseline activity before the onset of the stimulus), transient segment (first 200 milliseconds after the onset of the stimulus) and, sustained segment (200-1000 milliseconds after the onset of the stimulus). The sustained segments of evoked responses provided the most stable measures for analysis; Hence, they were used for subsequent analyses to quantify the effect of TMS.

In order to determine the effect of TMS on neuronal populations, it was important to first ascertain which channels (out of 96 implanted channels) demonstrated orientation tuning. This was achieved by computing the circular variance of direction tuning curves (Ringach et al., 1997; Ringach et al., 2002). In mathematical terms, circular variance (CV) for a bi-modal distribution, such as that expected for direction selectivity (Shmuel and Grinvald, 1996; Swindale et al., 2003), is given by

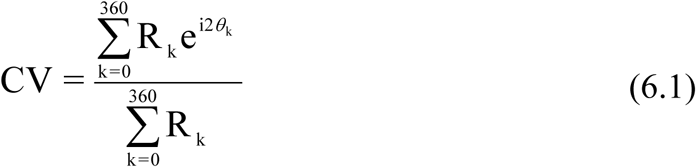

where, R_k_ is the response – firing rate – corresponding to direction *k* of the gratings and θ_k_ represents the direction angle of the drift of the gratings. The CV can be further simplified for uni-modal distribution, such as that expected for orientation selectivity (Hubel and Wiesel, 1962)

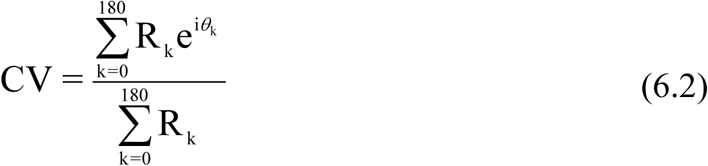

To weed out channels that showed tuning from channels that did not, we calculated the CV obtained for each of the 96 channels based on the action potential responses. A CV threshold of 0.9 was selected for separating tuned and un-tuned channels.

## Results

### The effect of TMS on firing rates and gamma band power

To quantify the effect of TMS right after its delivery, we analysed the data from the channels that were tuned to orientation. This was achieved by plotting the time courses of firing rates (FRs) and band-limited magnitude of gamma activity (gBLM). Figure 3 presents the time courses of the averaged FRs, low gBLM (30-57 Hz) and high gBLM (63-150 Hz) for each rTMS paradigm. Each individual measure (FR or low/high gBLM) was first normalized separately for each channel and condition by its just-pre-TMS value (5^th^ time point on the plots). The normalized responses were then averaged across all orientations, tuned channels and, finally across all hemispheres. Both cTBS and 15 Hz rTMS paradigms resulted in considerable suppression of spiking activity right after their administration. cTBS and 15 Hz rTMS suppressed spiking activity for 6.8 and 1.7 minutes, respectively (t-test, p<0.05, FDR corrected, N = 8; represented by blue ‘*’ markers near the lower y-axis extremity). In contrast, iTBS showed no suppressive effect after its delivery. iTBS caused statistically significant but sporadic long term facilitation in spiking activity (represented by red ‘*’ markers near the upper y-axis extremity). The changes were less obvious In low and high gBLM. However, the trends were similar: cTBS and 15 Hz rTMS suppressed gamma-band activity for 1.7 minutes after their administration with no significant long term effects. In contrast, iTBS showed no short term effects but a significant increase in long term gBLM, which was more consistent in high gBLM than in low gBLM.

**Figure 3.**
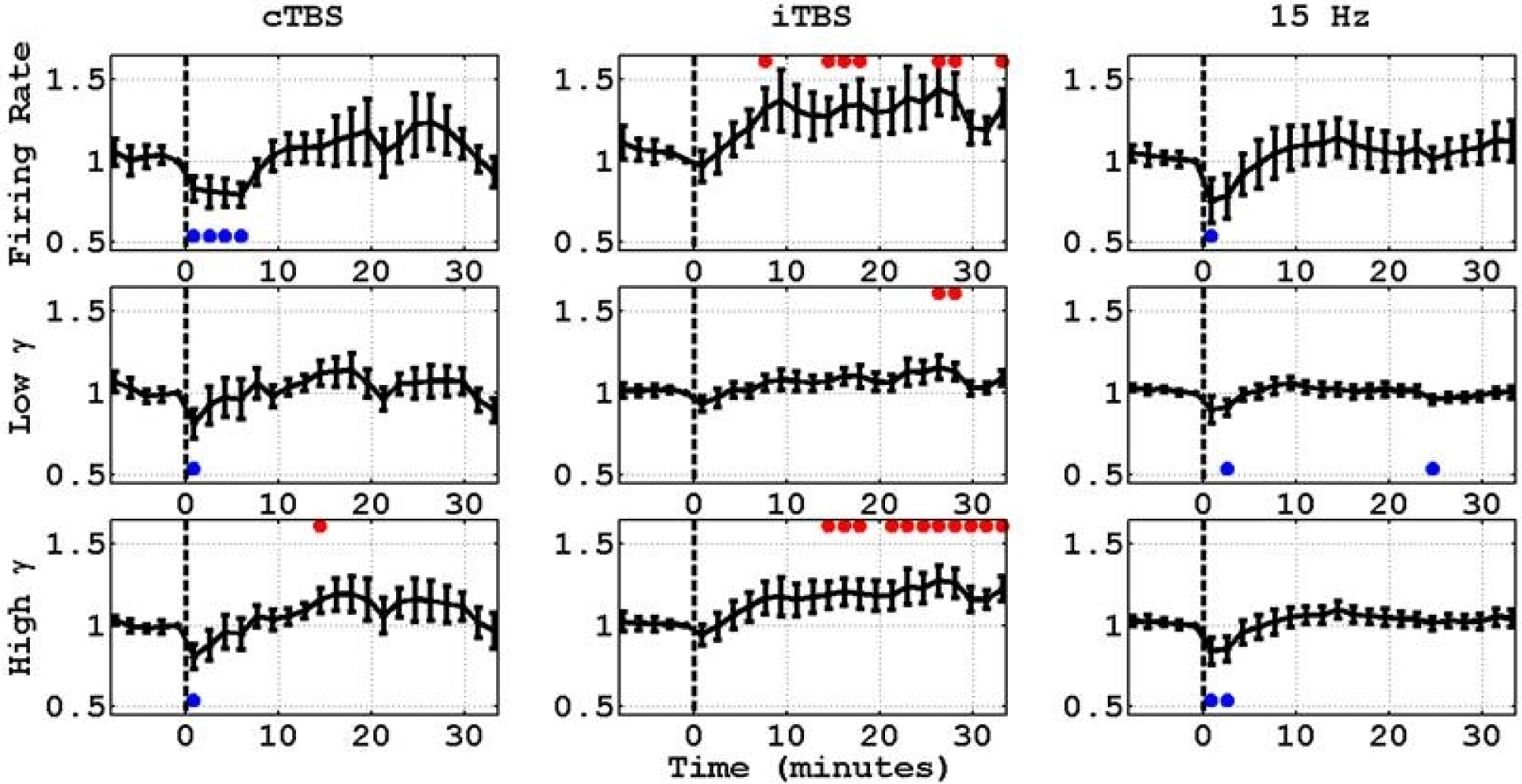
Average normalized action potentials firing rate and gBLM responses in cTBS, iTBS, and 15 Hz rTMS experiments. In all cases, normalization was done with respect to responses on the time point right before the application of TMS.

### The effect of TMS on the spatial pattern of the response

In the previous section, we examined the effects of rTMS on the average firing rate and gamma activity across all orientations and tuned channels. How these effects influence the spatial pattern of the responses to each orientation? To address this question, we compared the post-rTMS responses recorded in all channels to all orientations relative to their pre-rTMS counterparts. The comparison was done with reference to the pattern of responses recorded in the just-pre-TMS trial. This gave us a measure of how the response pattern of each trial varied with respect to the pre-TMS trial. To test a possible linear relationship between the pre- and post-TMS responses, we chose not to use simple linear regression, because simple regression assumes that one of the two variables – the one represented along the horizontal axis – is given with no errors. Here, both the pre-TMS and post-TMS responses are subject to noise and measurement errors. Therefore, we applied orthogonal regression, which assumes and models errors along the two axes (Leng et al., 2007). Figure 4 presents a snippet of the results, showing a scatter plot and the orthogonal regression line fits of post-TMS responses at different time points (1.7, 6.8 and 18.7 minutes after TMS) relative to the pre-TMS responses to all orientations averaged over the 3 trials (1.7 minutes) before TMS. The scatter plots show an approximate linear behavior of the responses at different time points. The orthogonal regression line for each cluster depicts the major axis of anisotropy. The slope of each regression represents a multiplicative factor, the gain associated with that cluster relative to the pre-TMS baseline. For the given experiments presented in Figure 4 – one each for cTBS, iTBS and 15 Hz rTMS conditioning selected randomly – we observe that cTBS conditioning resulted in a decrease in spiking activity after 2 minutes and 7 minutes (slopes of the red and green regression lines vs. the slope of the black unity line); Nineteen minutes following the application of cTBS, the reponses already returned to baseline (slope of blue regression line vs. slope of black unity line). iTBS conditioning resulted in a prolonged increase in cortical activity (shown by the higher slope of the blue regression line with respect to the unity line); this effect was not visible immediately following TMS administration (as indicated by the overlap of the red and black regression lines). 15 Hz rTMS resulted in a decrease in cortical responses following the delivery of TMS; Nineteen minutes following the application of 15 Hz TMS, the responses showed a slight increase relative to the baseline. These results demonstrated a linear gain effect of the three rTMS paradigms we applied, albeit from only one experiment and only 3 time points.

**Figure 4.**
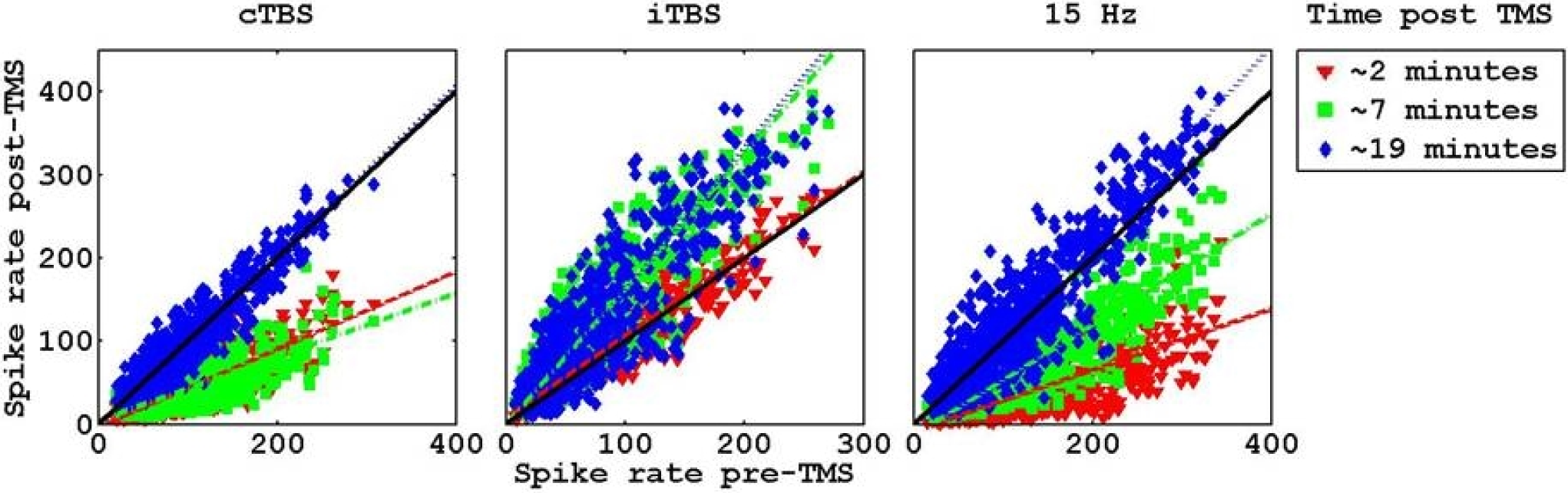
Linear gain effect of TMS: Post-Vs Pre TMS responses to oriented grating stimuli for cTBS, iTBS, and 15 Hz rTMS conditioning. Each plot presents the average spiking response recorded by tuned channels after the delivery of a particular rTMS conditioning paradigm with respect to average spiking response recorded during the sustained part of the response by the same tuned channels before the delivery of rTMS, for a specific experiment run. Averaging took place across the prolonged part of the response (200-1000 ms following the presentation of the stimulus) and over three consecutive trials, separately for each channel and orientation. Responses obtained 1.7, 6.8 and 18.7 minutes (portrayed by red, green and blue markers, respectively) after the delivery of the particular rTMS paradigm were compared to responses averaged over 1.7 minutes before TMS delivery. The colored scatter plots represent the relationship between post-TMS responses to the pre-TMS response. The corresponding colored lines for the scatter plots are the orthogonal regression lines. The black lines represent the lines with slope 1 and intercept 0 (unity slope lines).

Our subsequent analysis meant to test whether the effects of the three rTMS paradigms can be modelled as linear gains for the entire data-set. Figure 5 presents the time-courses of the slopes and y-intercepts of the regression lines along with the correlation coefficients. The three parameters represent global measures of neuronal responses since they represent changes across channels and orientations as a function of time. The regression slope represents a multiplicative factor by which responses were either suppressed or facilitated relative to the pre-rTMS response. The y-intercept refers to an additive component of responses relative to the pre-TMS baseline responses. The correlation coefficients indicate the similarity of the patterns of responses relative to the pre-rTMS pattern of responses. Each of these time courses of parameters was obtained separately for each experiment. Figure 5 presents the time-course averaged separately across all cTBS, iTBS and 15 Hz runs. The lower row of panels in Figure 5 presents the time courses of the correlation between each response pattern and the just-pre-TMS response pattern, averaged across experiments.

**Figure 5.**
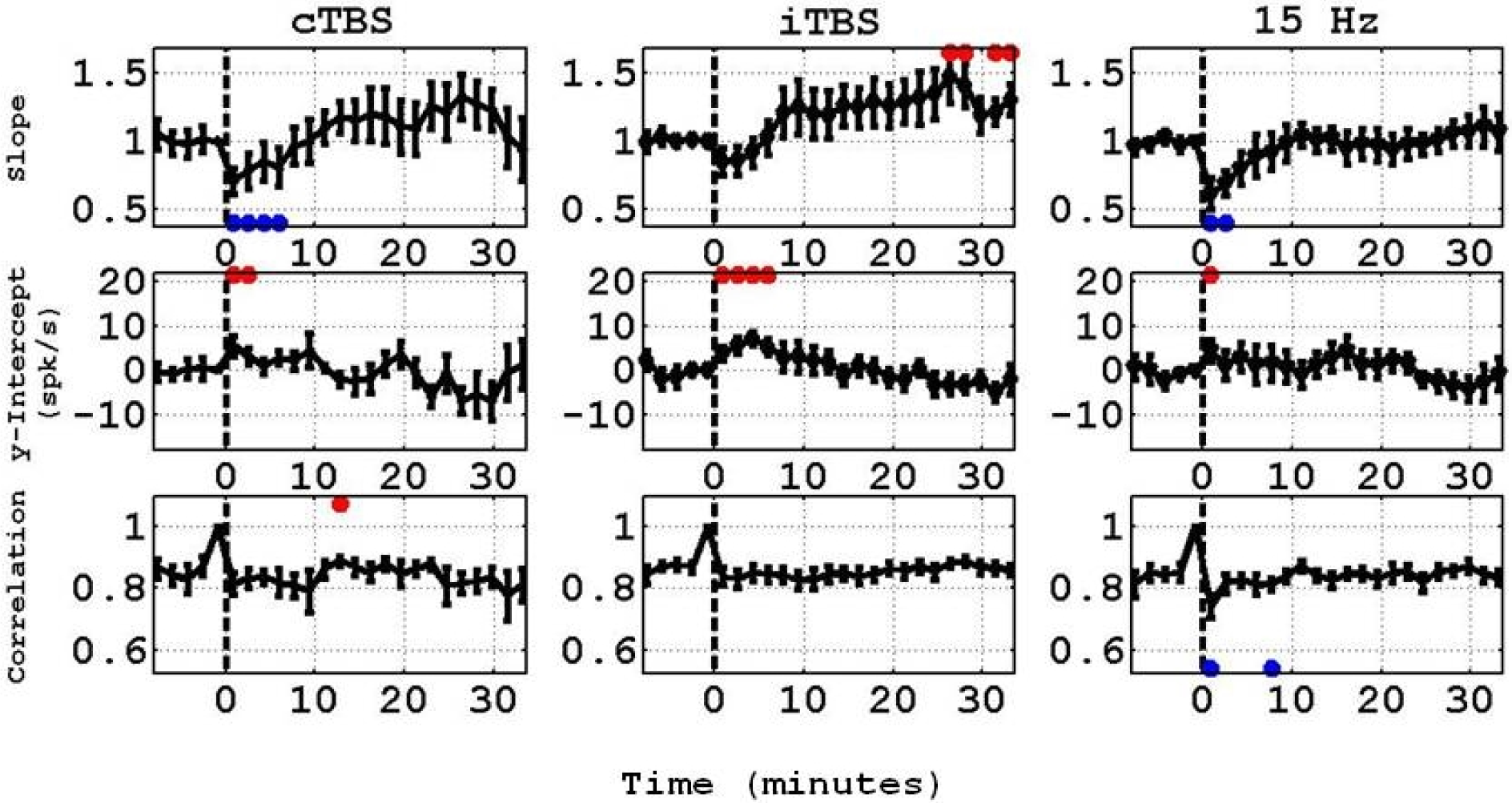
Time courses of the slopes and y-intercepts of the orthogonal regression lines and correlation coefficients applied to the scatter plots. The slope and intercept time courses represent the time point by time point results of orthogonal regression of responses to oriented gratings relative to the baseline responses recorded pre-TMS, for cTBS, iTBS and 15 Hz rTMS. The error bars represent the standard errors of mean (N = 8 experiments).

Table I presents the values of mean correlation coefficient values pre-rTMS (time points 1 to 4 or trials 1 through 12) and immediately after (time points 6 to 9 or trials 16 through 27) the application of rTMS. Based on these values we conclude that the application of TMS does not significantly change the response patterns (except for a 1.7 minutes-long transient effect right after the application of 15 Hz TMS).

**Table I.** Variation of the mean correlation coefficient values across trials before and after the delivery of rTMS (cTBS, iTBS, and 15 Hz rTMS conditioning).

| **Correlation**<br>**rTMS Paradigm** | ***Pre-TMS trials (1 to 12)***<br>***( Mean ±SEM)*** | ***Post-TMS trials (16 to 27)***<br>***(Mean ±SEM)*** |
| --- | --- | --- |
| *cTBS* | 0.853 ± 0.035 | 0.826 ± 0.034 |
| *iTBS* | 0.865 ± 0.020 | 0.841 ± 0.028 |
| *15 Hz* | 0.840 ± 0.029 | 0.799 ± 0.030 |

Mean correlation coefficients before (time points 1 to 4 or trials 1 through 12) and immediately after (time points 6 to 9 or trials 16 through 27) the application of rTMS. Each of the response patterns was compared to the response pattern from time point 5 (trials 13-15), which served as the pre-rTMS baseline response. The table reports mean correlation coefficient ± standard error of means (SEM) computed over the correlation coefficients obtained from 8 hemispheres.

In summary, in all 3 paradigms, the effect of rTMS on spiking activity response is approximately linear relative to pre-TMS responses, showing short term suppression in cTBS and 15Hz, and long term facilitation in iTBS (and a non-significant trend of facilitation in cTBS). However, the response pattern remains unchanged.

### The effect of TMS on orientation selectivity and tuning

The cortical sites that were classified as tuned to orientation are expected to show different firing rate responses to different orientations. If the effect of TMS is indeed linear relative to responses prior to TMS, we should expect that the orientation tuning will not change following the application of TMS. To test this expectation directly, we analyzed orientation tuning before and after TMS. Orientation tuning curves were obtained by first averaging the responses to the two opposite directions, both orthogonal to the same orientation. Therefore, the bi-modal direction tuning curves were transformed to unimodal orientation tuning curves spanning 0 to 180 degrees. The tuning curves were fitted with Von-Mises function (Swindale, 1998; Swindale et al., 2003) to which we added a variable to account for an additive offset (‘DC’):

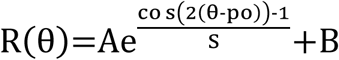

where R is the response to gratings of orientation θ°, A is the response at the preferred orientation, B is the DC offset, po is the preferred orientation and S is the spread parameter. The motive to add a DC parameter was to account for spontaneous activity and untuned visual evoked response analogous to a DC offset. In other words, the global signal (untuned evoked responses) was expected to be fitted by the DC offset parameter.

We first determined whether TMS changed the preferred orientation associated with each electrode. The upper row in Figure 6 presents the preferred orientation of all tuned channels in one experiment, obtained from the fitted tuning curves. The curves show that barring noise, there was no systematic change in preferred orientation as a function of time. In order to quantify the effect of TMS on the preferred orientation, we computed the difference in preferred orientation for each channel relative to the channel’s preferred orientation right before the administration of rTMS. The distributions in the second row in Figure 6 depict the preferred orientation at the beginning of the TMS session (time point 1, 6.8-8.5 minutes pre-TMS) and after the TMS conditioning (time point 9, 6.8-8.5 minutes post-TMS) relative to time point 5, just before TMS). These distributions are cumulative, including data from all tuned channels from all experiments. There was no statistically significant difference between the two distributions (D = 0.064, 0.057 and 0.039; χ^2^ = 5.95, 21.52, 18.72 where D is the Kolmogorov-Smirnov test statistic and χ^2^ is the Chi Square goodness of fit statistic for the distributions associated with cTBS, iTBS and 15 Hz rTMS, respectively). Based on the similarity of the distributions before and after TMS, we concluded that TMS did not change the preferred orientation.

**Figure 6.**
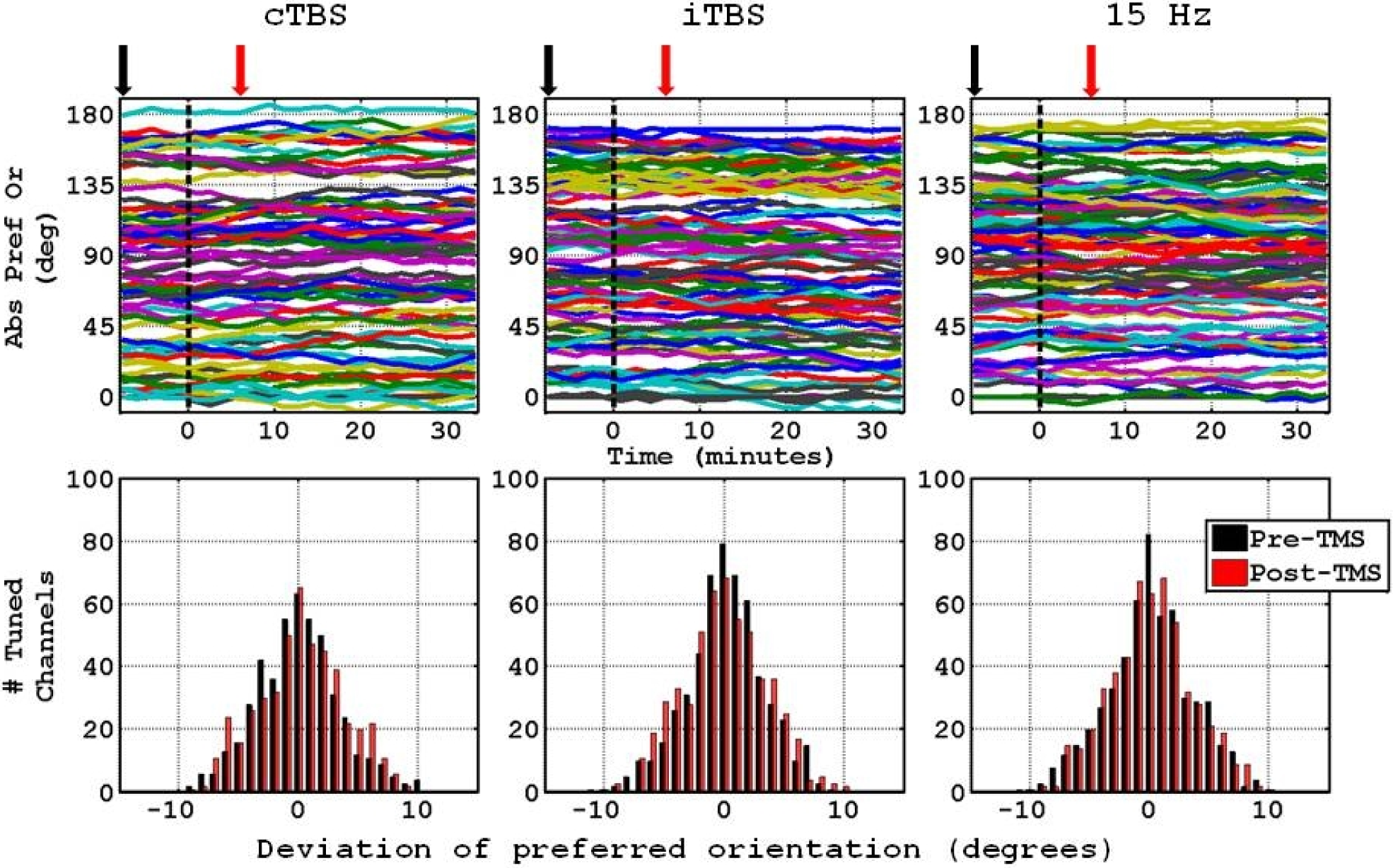
The upper panels present time courses of preferred orientation of each tuned channel, from one experiment with cTBS, iTBS, and 15 Hz rTMS. The lower panels present cumulative distributions of the maximum deviation of the preferred orientations of all tuned channels across all experiments. The maximum deviation is computed relative to the preferred orientations right before TMS, i.e. time point 5). In black, the deviations recorded at the beginning of the experiment session (−6.8 to −8.5 minutes with respect to when TMS is applied, also denoted by the black arrows in the upper panels). In red, the deviations recorded following the administration of rTMS (6.8 to 8.5 minutes following rTMS, also denoted by the red arrows in the upper panels).

In order to analyze the effect of TMS on other parameters of orientation tuning, we aligned all fitted tuning curves according to their preferred orientation such that the preferred orientation corresponded to 90°. Figure 7 shows plots of the average tuning curves fitted to the average of three trials right before TMS (in dark) and to the average of three trials right after TMS (in red). These tuning curves were averaged across all tuned channels and all experiments. The curves demonstrate the suppressive effect of cTBS and 15 Hz right after TMS, and the weak suppression following iTBS. Note that the relative change in firing rate for a given orientation following TMS relative to the average firing rate for the same orientation prior to TMS seems to remain approximately equal across all orientations. This means that the responses to the oriented gratings post-TMS bore a linear relationship with the response responses pre-TMS. An additional support to this observation is demonstrated by the vertical lines representing the full-width at half height before and after TMS, which are approximately equal.

**Figure 7.**
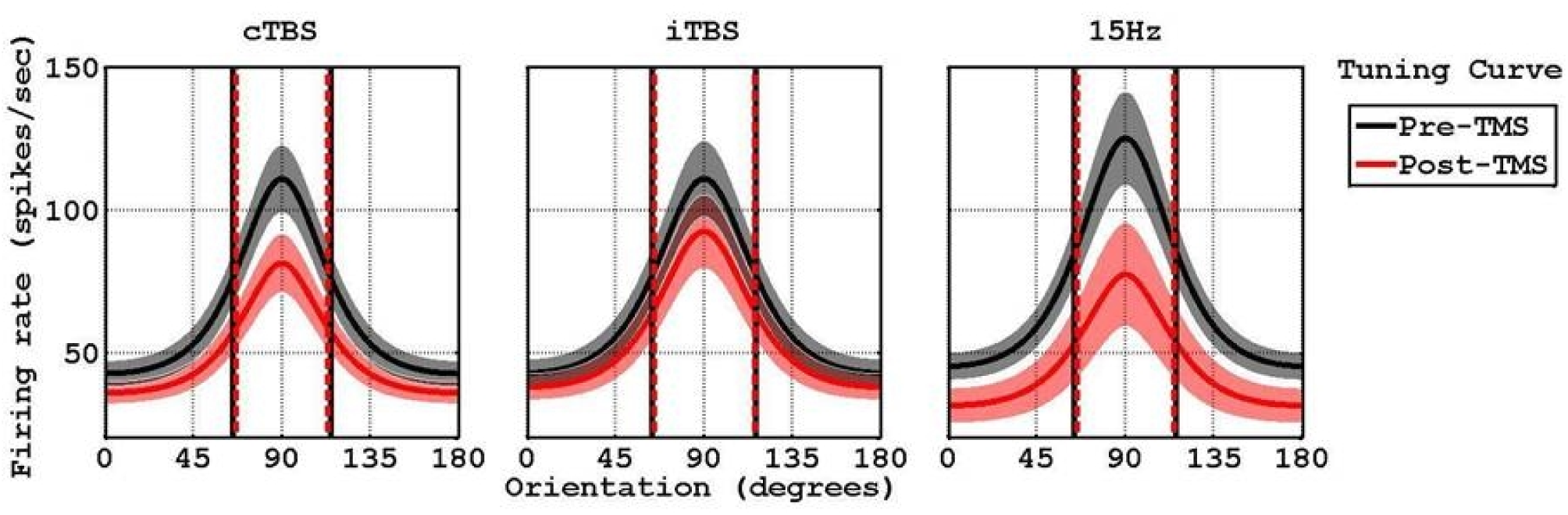
Average tuning curves of three trials right before (black) and right after (red) the administration of cTBS, iTBS, and 15 Hz rTMS. The darker lines represent the mean values whereas the lighter regions around them show the standard error of means across all experiments. The vertical solid black and dotted red lines mark the widths of the tuning curves at the corresponding half heights before and after TMS, respectively.

Figure 8 presents the time courses of the average tuning curves for the three TMS paradigms. The black and red arrows indicate the time points at which the tuning curves shown in Figure 7 were extracted from.

**Figure 8.**
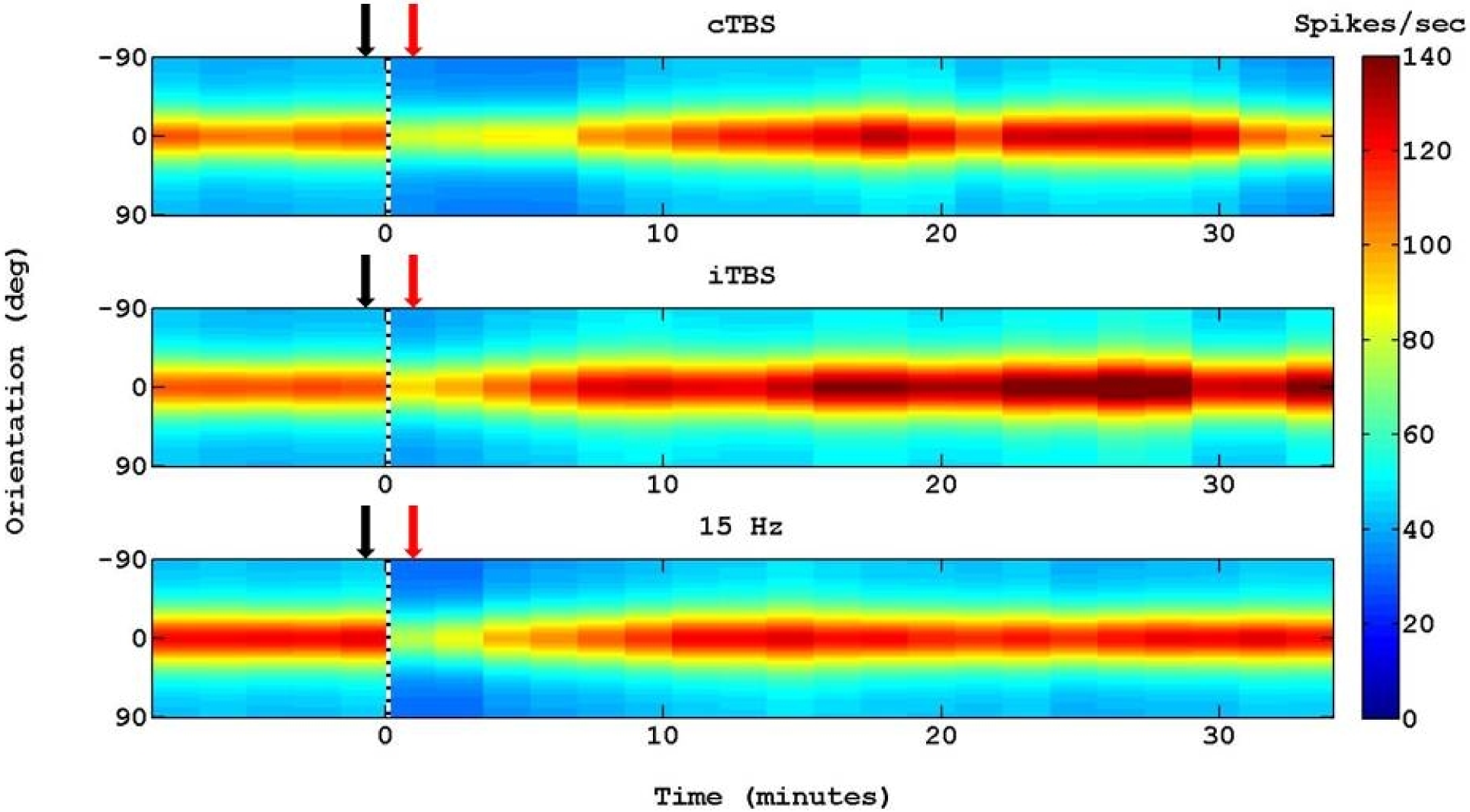
Time-courses of average tuning curves fitted with Von-Mises function. The dotted black vertical lines separate the tuning curves into the pre-TMS and post-TMS regions. The tuning curve of spike firing rate obtaind from each recording site was fitted with Von-Mises function. Each of the fitted function curves was circularly shifted such that the preferred orientations of all fitted curves were aligned before averaging across all recording sites.

Figure 9 presents the time courses of 1) the mean relative difference in firing rate response amplitude between the preferred and non-preferred orientation (orientation that elicits the lowest firing rate). Note that ‘relative’ refers to that same measure in time point 5, just before TMS; 2) non-preferred orientation firing rate (DC parameter); 3) half-width at half maximum height (HW@HM), also termed the bandwidth of the tuning curve; and, 4) circular variance (CV). CV is a global measure of tuning and selectivity whereas the bandwidth is a local measure (Ringach et al., 2002). Consistent with the results presented in Figure 3 and Figure 5, cTBS and 15 Hz TMS suppressed the amplitude of the response of the tuning curve for 6.8 and 1.7-3.4 minutes, respectively. Consistent with the results presented in Figures 3-5, iTBS caused a late (14-32 min following TMS) increase in tuning curve amplitude. TMS caused a transient (1.7 min long) increase in CV right after cTBS conditioning. No statistically significant changes in tuning bandwidth were observed following each of the 3 TMS paradigms. We concluded that none of the TMS paradigms affected orientation tuning and selectivity. We further concluded that the effect of TMS on the response amplitude is linear relative to the baseline pre-TMS response.

**Figure 9.**
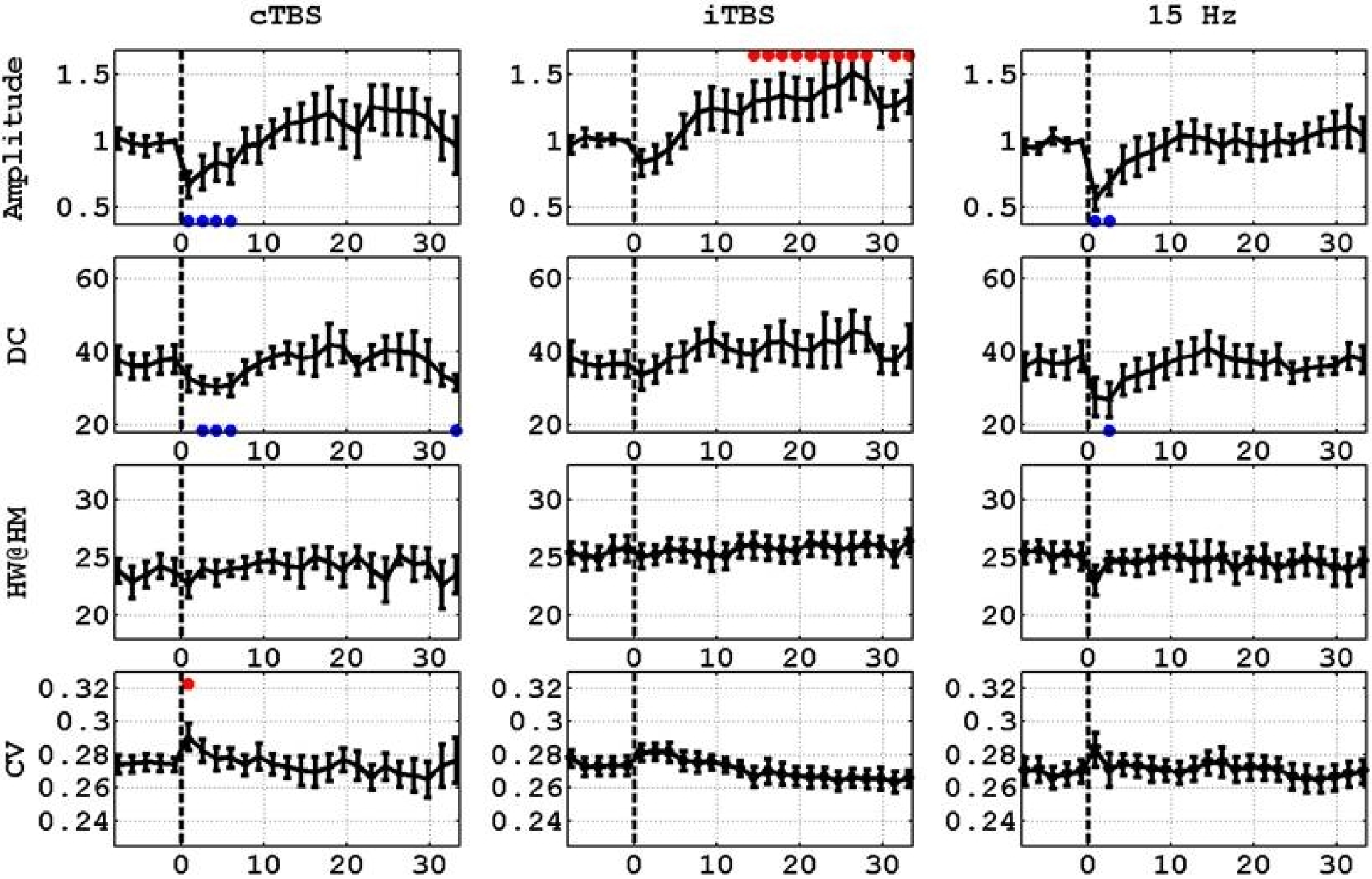
Time courses of the amplitude (response to preferred orientation minus the response to the orthogonal-to-preferred orientation, normalized to the same measure obtained just-prior to TMS), DC (response to the non-preferred orientation) half-width at half-maximum-height and circular variance of the fitted tuning curves. The left, middle and right columns present the values obtained from applying cTBS, iTBS, and 15 Hz rTMS, respectively.

## Discussion

Simulations show that the current density elicited by TMS is approximately homogeneous in the region covered by the multi-electrode array (Romero et al., 2019). A standard 70-mm butterfly coil – which we used in our study – is thought to maximally stimulate about 1–2 cm^2^ of cortex beneath its central junction (Sandrini et al., 2011). Therefore, the effect of TMS spreads out over a larger area than that covered by the electrode array.

Is the effect of TMS on the neuronal assemblies within the region influenced by the TMS similar? Is it additive, multiplicative or non-linear for different populations of neurons within that area? Or are neuronal assemblies, be it strongly responding or weakly responding, have their performance after the application of TMS affected by the same percentage relative to their responses before the application of TMS? Is the effect exerted by different rTMS protocols similar? Here we addressed these questions by applying different rTMS protocols on a 3.6 mm × 3.6 mm area of the feline visual cortex.

### Effect of rTMS on neuronal responses

Based on previous studies in humans, low-frequency and high-frequency rTMS decreases and increases cortical excitability of the motor cortex, respectively (Berardelli et al., 1998; Fitzgerald et al., 2006).

Data from animal studies using TMS in combination with direct electrophysiological recording have provided insight into the TMS effects. Moliadze et al. (2003) performed extracellular single-unit recordings in the cat primary visual cortex (area 17) while directing a 70 mm figure of eight coil – similar to the one we used – to the recording site. Consistent with the findings on low-frequency TMS in humans, single biphasic TMS pulses elicited facilitation of activity during the first 500 ms, followed thereafter by a suppression of activity lasting up to a few seconds.

In contrast to the agreement on the effect of low-frequency TMS, animal studies that applied high-frequency TMS reported suppression of neuronal activity, unlike the findings from the human motor cortex. Valero-Cabre et al. (2005) demonstrated that the application of 20 Hz rTMS induces a relative decrease in metabolic activity at the site of direct application and at specific distant sites of the stimulated hemisphere, indicating an rTMS-induced decrease in neural activity. Allen et al. (2007) and Pasley et al. (2009) applied TMS to the cat visual cortex and evaluated the consequences to neuronal activity. In contrast to findings observed in the human motor cortex, short TMS pulse trains (1 to 4 s, 1 to 8 Hz) elicited an immediate suppression of action potential and LFP responses that lasted 5 to 10 minutes (Allen et al., 2007; Pasley et al., 2009). The immediate, short suppression effect we observed after applying 15 Hz rTMS (Figures 3-5 and 7-9) is consistent with the response suppression elicited by Moliadze et al. (2003) following the application of single pulses, and that observed by Allen et al. (2007) following the application of 8 Hz TMS. The differences in the effect durations observed in the different studies – Moliadze et al. (2003), Allen et al. (2007), and our study (Figures 3-5 and 7-9) can be attributed to the different frequencies applied in the 3 studies, and to the different durations of rTMS.

Huang et al. (2005) were the first to introduce TBS in humans. Application of 600 pulses of cTBS reduces the excitability of the motor cortex, as indicated by the motor evoked potentials measured in a hand muscle (Huang et al., 2005). In contrast, iTBS increased this excitability, and 15 Hz rTMS did not show significant effects. Our intra-cortical measurements are – by large – aligned with the effects of cTBS and iTBS on the motor cortex in humans (Huang et al., 2005). Analogous parameters recorded from different regions of the brain can be affected differently by the same TMS paradigm. We hypothesize that the differences between the findings from the human motor cortex by Huang et al. (2005) and our findings from the cat visual cortex may be attributed to the different laminar structure of the motor and visual cortex..

### TMS does not influence the global pattern of responses

Linearity and correlation analysis of firing rates revealed that TMS influences the responses of neuronal assemblies in orientation columns the same way. In other words, the response of multi-unit activity across all orientations – be it preferred or non-preferred orientation – bears a linear gain effect relationship to their corresponding responses before TMS, with the same multiplicative factor across recordings from all electrodes. This was also demonstrated using tuning curve analysis in which we show that TMS did not affect the preferred orientation and the global measures of tuning and selectivity (Figures 6-9). These findings indicate that the rTMS paradigms do not change the functionality of the circuit. Rather, they modify the amplitude of the responses.

In conclusion, TMS effects on cat area 18 responses to grating stimuli can be modeled as linear gain effects. This can be explained by the current density elicited with a figure-of-eight coil, which is approximately constant in the 3.6 mm × 3.6 mm region where the electrode array is implanted. The state of ensembles of neurons in a population within a constant field distribution is affected to the same extent by TMS.

### Neuronal mechanisms of rTMS

The precise mechanisms of TMS action are still unclear. Variuos hypotheses have been made with regard to the neuronal mechanisms underlying the effect of TMS and rTMS.

Simultaneous recording of LFPs and spiking activity in the primary visual cortex of macaque monkeys showed that the spatial extent of gamma activity and its relationship to local spiking activity is stimulus-dependent (Jia et al., 2011). Small gratings induce a broadband increase in spectral power. Large gratings induce a gamma rhythm characterized by a distinctive spectral “bump,” which is coherent across widely separated sites. These results are consistent with the increasing amplitude of the intrinsic signal responses with increasing anisotropy of grating stimuli (Shmuel et al., 1996). Previous studies showed that TMS disrupts the temporal structure of activity by altering phase relationships between neural signals (Allen et al., 2007; Pasley et al., 2009), specifically between neurons belonging to a more extended circuit activated by the stimulus (Pasley et al., 2009; Sandrini et al., 2011). This could well be a mechanism that contributes to our findings too. Indeed, a possible disruption of the high coherence elicited by grating stimuli is a plausible mechanism underlying the suppression effects cTBS and 15 Hz TMS elicited in our study.

A plausible hypothesis is that TMS induces intracortical inhibition (Sandrini et al., 2011). Indeed, an induced electric field can increase GABA levels (Dubin et al., 2016), which in turn, suppresses activity. Our finding of a linear gain effect suppression of neuronal responses following cTBS and 15 Hz TMS, with no changes observed to the preferred orientation or the tuning of orientation preference is consistent with elevated levels of GABA.

## Acknowledgments

This work was supported by a grant from the Natural Sciences and Engineering Research Council of Canada and the Canadian Institute of Health Research (CHRP 385962-10). We thank Drs. Lisa Koski, Frederic Lesage, and Christophe Grova for their very helpful comments.

## References

Allen, E.A., Pasley, B.N., Duong, T., Freeman, R.D., 2007. Transcranial magnetic stimulation elicits coupled neural and hemodynamic consequences. Science 317, 1918–1921.

Awad, B.I., Carmody, M.A., Zhang, X., Lin, V.W., Steinmetz, M.P., 2013. Transcranial Magnetic Stimulation After Spinal Cord Injury. World Neurosurgery.

Baker, S.N., Olivier, E., Lemon, R.N., 1995. Task-related variation in corticospinal output evoked by transcranial magnetic stimulation in the macaque monkey. The Journal of Physiology 488, 795–801.

Barker, A.T., Freeston, I.L., Jalinous, R., Jarratt, J.A., 1985a. Non-invasive stimulation of motor pathways within the brain using time-varying magnetic fields. Electroencephalography and Clinical Neurophysiology 61, S245–S246.

Benali, A., Trippe, J., Weiler, E., Mix, A., Petrasch-Parwez, E., Girzalsky, W., Eysel, U.T., Erdmann, R., Funke, K., 2011. Theta-Burst Transcranial Magnetic Stimulation Alters Cortical Inhibition. The Journal of Neuroscience 31, 1193–1203.

Berardelli, A., Inghilleri, M., Rothwell, J.C., Romeo, S., Currà, A., Gilio, F., Modugno, N., Manfredi, M., 1998. Facilitation of muscle evoked responses after repetitive cortical stimulation in man. Experimental Brain Research 122, 79–84.

Berlim, M.T., Van den Eynde, F., Daskalakis, Z.J., 2012. A systematic review and meta-analysis on the efficacy and acceptability of bilateral repetitive transcranial magnetic stimulation (rTMS) for treating major depression. Psychological Medicine FirstView, 1–10.

Berlim, M.T., Van den Eynde, F., Daskalakis, Z.J., 2013. Efficacy and acceptability of high frequency repetitive transcranial magnetic stimulation (rTMS) versus electroconvulsive therapy (ECT) for major depression: A systematic review and meta-analysis of randomized trials. Depression and Anxiety.

Brainard, D.H., 1997. The Psychophysics Toolbox. Spatial Vision 10, 433–436.

Dancause, N., Barbay, S., Frost, S.B., Zoubina, E.V., Plautz, E.J., Mahnken, J.D., Nudo, R.J., 2006. Effects of Small Ischemic Lesions in the Primary Motor Cortex on Neurophysiological Organization in Ventral Premotor Cortex. Journal of Neurophysiology 96, 3506–3511.

de Labra, C., Rivadulla, C., Grieve, K., Mariño, J., Espinosa, N., Cudeiro, J., 2007. Changes in Visual Responses in the Feline dLGN: Selective Thalamic Suppression Induced by Transcranial Magnetic Stimulation of V1. Cerebral Cortex 17, 1376–1385.

De Ridder, D., van der Loo, E., Van der Kelen, K., Menovsky, T., van de Heyning, P., Moller, A., 2007. Theta, alpha and beta burst transcranial magnetic stimulation: brain modulation in tinnitus. International journal of medical sciences 4, 237–241.

Di Lazzaro, V., Pilato, F., Dileone, M., Profice, P., Oliviero, A., Mazzone, P., Insola, A., Ranieri, F., Meglio, M., Tonali, P.A., Rothwell, J.C., 2008. The physiological basis of the effects of intermittent theta burst stimulation of the human motor cortex. The Journal of Physiology 586, 3871–3879.

Dubin, M.J., Mao, X., Banerjee, S., Goodman, Z., Lapidus, K.A.B, Kang, G., Liston, C., Shungu, D.C., 2016. Elevated prefrontal cortex GABA in patients with major depressive disorder after TMS treatment measured with proton magnetic resonance spectroscopy. Journal of Psychiatry and Neuroscience, 41:E37–E45.

Ernst, E., 1990. A review of stroke rehabilitation and physiotherapy. Stroke 21, 1081–1085.

Espinosa, N., de Labra, C., Rivadulla, C., Mariño, J., Grieve, K.L., Cudeiro, J., 2007. Effects on EEG of Low (1Hz) and High (15Hz) Frequency Repetitive Transcranial Magnetic Stimulation of the Visual Cortex: A Study in the Anesthetized Cat. The Open Neuroscience Journal 1, 20–25.

Espinosa, N., Mariño, J., de Labra, C., Cudeiro, J., 2011. Cortical Modulation of the Transient Visual Response at Thalamic Level: A TMS Study. PloS one 6, 1–11.

Gersner, R., Kravetz, E., Feil, J., Pell, G., Zangen, A., 2011. Long-Term Effects of Repetitive Transcranial Magnetic Stimulation on Markers for Neuroplasticity: Differential Outcomes in Anesthetized and Awake Animals. The Journal of Neuroscience 31, 7521–7526.

Grinvald, A., Lieke, E., Frostig, R.D., Gilbert, C.D., Wiesel, T.N., 1986. Functional architecture of cortex revealed by optical imaging of intrinsic signals. Nature 324, 361–364.

Huang, Y.-Z., Edwards, M.J., Rounis, E., Bhatia, K.P., Rothwell, J.C., 2005. Theta Burst Stimulation of the Human Motor Cortex. Neuron 45, 201–206.

Hubel, D.H., Wiesel, T.N., 1962. Receptive fields, binocular interaction and functional architecture in the cat’s visual cortex. J Physiol 160, 106–154.

Ishikawa, S., Matsunaga, K., Nakanishi, R., Kawahira, K., Murayama, N., Tsuji, S., Huang, Y.-Z., Rothwell, J.C., 2007. Effect of theta burst stimulation over the human sensorimotor cortex on motor and somatosensory evoked potentials. Clinical Neurophysiology 118, 1033–1043.

Jia X., Matthew A., Smith, M.A., Kohn, A., 2011. Stimulus Selectivity and Spatial Coherence of Gamma Components of the Local Field Potential. Journal of Neuroscience, 31, 9390–9403; DOI: https://doi.org/10.1523/JNEUROSCI.0645-11.2011

Joo, E.Y., Han, S.J., Chung, S.-H., Cho, J.-W., Seo, D.W., Hong, S.B., 2007. Antiepileptic effects of low-frequency repetitive transcranial magnetic stimulation by different stimulation durations and locations. Clinical Neurophysiology 118, 702–708.

Khedr, E.M., Abdel-Fadeil, M.R., Farghali, A., Qaid, M., 2009. Role of 1 and 3 Hz repetitive transcranial magnetic stimulation on motor function recovery after acute ischaemic stroke. European Journal of Neurology 16, 1323–1330.

Leng, L., Zhang, T., Kleinman, L., Zhu, W., 2007. Ordinary least square regression, orthogonal regression, geometric mean regression and their applications in aerosol science. Journal of Physics: Conference Series 78, 012084.

Leong, K.W., 2009. Predicting the response to neural stimulation in rehabilitation of stroke-related motor impairment. McGill University, Montreal.

Levkovitz, Y., Marx, J., Grisaru, N., Segal, M., 1999. Long-Term Effects of Transcranial Magnetic Stimulation on Hippocampal Reactivity to Afferent Stimulation. The Journal of Neuroscience 19, 3198–3203.

Maeda, F., Keenan, J.P., Tormos, J.M., Topka, H., Pascual-Leone, A., 2000a. Interindividual variability of the modulatory effects of repetitive transcranial magnetic stimulation on cortical excitability. Experimental Brain Research 133, 425–430.

Massé-Alarie, H., Flamand, V.H., Moffet, H., Schneider, C., 2013. Peripheral Neurostimulation and Specific Motor Training of Deep Abdominal Muscles Improve Posturomotor Control in Chronic Low Back Pain. The Clinical Journal of

Moliadze, V., Giannikopoulos, D., Eysel, U.T., Funke, K., 2005. Paired-pulse transcranial magnetic stimulation protocol applied to visual cortex of anaesthetized cat: effects on visually evoked single-unit activity. The Journal of Physiology 566, 955–965.

Moliadze, V., Zhao, Y., Eysel, U., Funke, K., 2003. Effect of transcranial magnetic stimulation on single-unit activity in the cat primary visual cortex. The Journal of Physiology 553, 665–679.

Noh, N.A., Fuggetta, G., Manganotti, P., Fiaschi, A., 2012. Long Lasting Modulation of Cortical Oscillations after Continuous Theta Burst Transcranial Magnetic Stimulation. PloS one 7, e35080.

Pascual-Leone, A., Davey, N.J., Rothwell, J., Wassermann, E.M., Puri, B.K., 2002. Handbook of Transcranial Magnetic Stimulation. Oxford University Press, London; New York, NY.

Pasley, B.N., Allen, E.A., Freeman, R.D., 2009. State-dependent variability of neuronal responses to transcranial magnetic stimulation of the visual cortex. Neuron 62, 291–303.

Prikryl, R., Kucerova, H., 2005. Occurrence of epileptic paroxysm during repetitive transcranial magnetic stimulation treatment. Journal of Psychopharmacology 19, NP.

Prikryl, R., Kucerova, H.P., 2013. Can Repetitive Transcranial Magnetic Stimulation Be Considered Effective Treatment Option for Negative Symptoms of Schizophrenia? The Journal of ECT 29, 67-74 10.1097/YCT.1090b1013e318270295f.

Rajji, T.K., Rogasch, N.C., Daskalakis, Z.J., Fitzgerald, P.B., 2013. Neuroplasticity-Based Brain Stimulation Interventions in the Study and Treatment of Schizophrenia: A Review. Les interventions de stimulation cérébrale basée sur la neuroplasticité dans l’étude et le traitement de la schizophrénie: une revue. 58, 93–98.

Ringach, D.L., Hawken, M.J., Shapley, R., 1997. Dynamics of orientation tuning in macaque primary visual cortex. Nature 387, 281–284.

Ringach, D.L., Shapley, R.M., Hawken, M.J., 2002. Orientation Selectivity in Macaque V1: Diversity and Laminar Dependence. The Journal of Neuroscience 22, 5639–5651.

Romero M.C., Davare, M., Armendariz, M., Janssen, P., 2019. Neural effects of transcranial magnetic stimulation at the single-cell level. Nature Communications, 10:2642. doi: 10.1038/s41467-019-10638-7.

Rossi, S., Hallett, M., Rossini, P.M., Pascual-Leone, A., 2009. Safety, ethical considerations, and application guidelines for the use of transcranial magnetic stimulation in clinical practice and research. Clin Neurophysiol 120, 2008–2039.

Rossini, P.M., Rossini, L., Ferreri, F., 2010. Brain-Behavior Relations: Transcranial Magnetic Stimulation: A Review. Engineering in Medicine and Biology Magazine, IEEE 29, 84–96.

Sandrini, M., Umiltac, C., Rusconi, 2011. The use of transcranial magnetic stimulation in cognitive neuroscience: A new synthesis of methodological issues. Neuroscience and Biobehavioral Reviews, 35:516–536.

Schabrun, S.M., Jones, E., Kloster, J., Hodges, P.W., 2013. Temporal association between changes in primary sensory cortex and corticomotor output during muscle pain. Neuroscience 235, 159–164.

Shmuel, A., Grinvald, A., 1996. Functional Organization for Direction of Motion and Its Relationship to Orientation Maps in Cat Area 18. The Journal of Neuroscience 16, 6945–6964.

Shmuel, A., Grinvald, A., 2000. Coexistence of linear zones and pinwheels within orientation maps in cat visual cortex. Proceedings of the National Academy of Sciences of the United States of America 97, 5568–5573.

Siebner, H.R., 2010. Can we enhance training-induced plasticity by modulating inhibitory cortical circuits with transcranial stimulation? (Commentary on Mix et al.). European Journal of Neuroscience 32, 1573–1574.

Swindale, N.V., 1998. Orientation tuning curves: empirical description and estimation of parameters. Biological Cybernetics 78, 45–56.

Swindale, N.V., Grinvald, A., Shmuel, A., 2003. The Spatial Pattern of Response Magnitude and Selectivity for Orientation and Direction in Cat Visual Cortex. Cerebral Cortex 13, 225–238.

Takeuchi, N., Chuma, T., Matsuo, Y., Watanabe, I., Ikoma, K., 2005. Repetitive Transcranial Magnetic Stimulation of Contralesional Primary Motor Cortex Improves Hand Function After Stroke. Stroke 36, 2681–2686.

Tergau, F., Naumann, U., Paulus, W., Steinhoff, B.J., 1999. Low-frequency repetitive transcranial magnetic stimulation improves intractable epilepsy. The Lancet 353, 2209.

Thielscher, A., Opitz, A., Windhoff, M., 2011. Impact of the gyral geometry on the electric field induced by transcranial magnetic stimulation. NeuroImage 54, 234–243.

Valero-Cabre, A., Payne, B.R., Pascual-Leone, A., 2007. Opposite impact on 14C-2-deoxyglucose brain metabolism following patterns of high and low frequency repetitive transcranial magnetic stimulation in the posterior parietal cortex. Exp Brain Res 176, 603–615.

Valero-Cabré, A., Payne, B.R., Rushmore, J., Lomber, S.G., Pascual-Leone, A., 2005. Impact of repetitive transcranial magnetic stimulation of the parietal cortex on metabolic brain activity: a 14C-2DG tracing study in the cat. Experimental brain research. Experimentelle Hirnforschung. Expérimentation cérébrale 163, 1–12.

Vernet, M., Bashir, S., Yoo, W.-K., Perez, J.M., Najib, U., Pascual-Leone, A., 2013. Insights on the neural basis of motor plasticity induced by theta burst stimulation from TMS–EEG. European Journal of Neuroscience 37, 598–606.

Vlachos, A., Müller-Dahlhaus, F., Rosskopp, J., Lenz, M., Ziemann, U., Deller, T., 2012. Repetitive Magnetic Stimulation Induces Functional and Structural Plasticity of Excitatory Postsynapses in Mouse Organotypic Hippocampal Slice Cultures. The Journal of Neuroscience 32, 17514–17523.

Wagner, T., Gangitano, M., Romero, R., Théoret, H., Kobayashi, M., Anschel, D., Ives, J., Cuffin, N., Schomer, D., Pascual-Leone, A., 2004a. Intracranial measurement of current densities induced by transcranial magnetic stimulation in the human brain. Neuroscience Letters 354, 91–94.

Wasserman, E.M., Zimmerman, T., 2012. Transcranial Magnetic Brain Stimulation: Therapeutic Promises and Scientific Gaps. Pharmacol Ther., 133: 98–107. doi: 10.1016/j.pharmthera.2011.09.003

Wu, T., Sommer, M., Tergau, F., Paulus, W., 2000. Lasting influence of repetitive transcranial magnetic stimulation on intracortical excitability in human subjects. Neuroscience Letters 287, 37–40.

Zapallow, C., Asmussen, M., Bolton, D.A., Lee, K.G., Jacobs, M., Nelson, A., 2012. Theta burst repetitive transcranial magnetic stimulation attenuates somatosensory evoked potentials from the lower limb. BMC Neuroscience 13, 133.

